# Surge of Corticocardiac Coupling in SHRSP Rats Exposed to Forebrain Cerebral Ischemia

**DOI:** 10.1101/388983

**Authors:** Fangyun Tian, Tiecheng Liu, Gang Xu, Talha Ghazi, Azeem Sajjad, Peter Farrehi, Michael M. Wang, Jimo Borjigin

## Abstract

Sudden death is an important but under-recognized consequence of stroke. Acute stroke can disturb central control of autonomic function, and result in cardiac dysfunction and sudden death. Previous study showed that bilateral common carotid artery ligation (BCCAL) in spontaneously hypertensive stroke-prone rats (SHRSP) is a well-established model for forebrain ischemic sudden death. This study aims to investigate the temporal dynamic changes in electrical activities of the brain and heart and functional interactions between the two vital organs following forebrain ischemia. EEG and ECG signals were simultaneously collected from 9 SHRSP and 8 Wistar-Kyoto (WKY) rats. RR interval and cardiac arrhythmias were analyzed to investigate the cardiac response to brain ischemia. EEG power and coherence (CCoh) analysis were conducted to study the cortical response. Corticocardiac coherence (CCCoh) and directional connectivity (CCCon) were analyzed to determine brain-heart connection. Heart rate variability (HRV) was analyzed to evaluate autonomic functionality. BCCAL resulted in 100% mortality in SHRSP within 14 hours, whereas no mortality was observed in WKY. The functionality of both the brain and the heart were significantly altered in SHRSP compared to WKY after BCCAL. SHRSP rats, but not WKY rats, exhibited intermittent surge of CCCoh, which paralleled the elevated CCCon and reduced HRV, following the onset of ischemia until sudden death. Elevated brain-heart coupling invariably associated with the disruption of the autonomic nervous system and the risk of sudden death. This study may improve our understanding of the mechanism of ischemic stroke-induced sudden death.

## Author Contributions

JB conceived the project and planned experiments. TL and FT conducted experiments. FT, GX, TG, and AS analyzed data. PF assisted with validation of cardiac arrhythmias. FT, MW, and JB wrote the paper.

## Introduction

Sudden death is an important but under-recognized consequence of stroke (Sörös and Hachinski, 2012). Despite major improvements in stroke diagnosis and treatment, 2-6% of patients suffer from sudden, unexpected death within the first 3 months after ischemic stroke (Prosser et al. 2007; Sörös and Hachinski 2012). In addition, about 19% of patients have fatal or serious non-fatal cardiac events, such as ventricular arrhythmias, which greatly increase the risk of sudden death (Prosser et al. 2007; Frangiskakis et al. 2009; Sörös and Hachinski 2012). The mechanisms of ischemic stroke-induced sudden death remain unclear. As a consequence, identification of patients at risk and prevention of stroke-induced sudden death present a major challenge.

Our laboratory has discovered that the brain is highly activated immediately following global ischemia induced by experimental cardiac arrest (Borjigin et al. 2013). More recently, we reported a marked surge of functional connectivity (cortical coherence, CCoh) and directional connectivity (cortical connectivity, CCon) in the dying brain of rats following asphyxia, which paralleled the excessive cortical release of a set of core neurotransmitters (Li et al. 2015a). In addition to the changes within the brain, a surge of corticocardiac coherence (CCCoh) and directional corticocardiac connectivity (CCCon), indicator for strong brain-heart connection, emerged after the onset of asphyxia with a reliable time delay (Li et al. 2015a). These data suggest that sudden death in rats that suffered global ischemia is tightly associated with the surge of functional connectivity within the cortex as well as between the brain and the heart. Whether this association exists in rats dying from forebrain ischemia is unknown.

Bilateral common carotid artery ligation (BCCAL) in stroke-prone spontaneously hypertensive rat (SHRSP) is a well-established model for forebrain ischemia (Kakihana et al. 1983; Lobanova et al. 2008). The cerebrovascular architecture and risk factors in SHRSP resemble with ischemic stroke in human patients, which makes SHRSP, derived from the normotensive Wistar-Kyoto (WKY) rat, a suitable model for studying ischemic stroke in rats (Yamori et al. 1976). Compared to asphyxic cardiac arrest rat model used in our previous studies, in which the entire body, including the brain and heart, is globally affected, the forebrain ischemia model (BCCAL) tests the direct influence of brain ischemia on cortical/cardiac functions, and permits the dissection of functional communication between the two vital organs. In this study, we use advanced signal processing techniques to functionally characterize the brain and the heart in the BCCAL model. Our goal is to understand how neurological injuries lead to abnormal brain-heart connection, autonomic dysfunction, cardiac damage, and sudden death. Ultimately, we hope this line of investigation will contribute to better understanding of the dying brain and the development of novel non-invasive biomarkers for prediction of risk for sudden death.

## Materials and methods

### Animals

Inbred SHRSP and WKY rats were acclimatized in our housing facility for at least 1 week before implantation of electrodes. Following electrode implantation, rats were allowed to recover for 1 week before online recording. The experimental procedures were approved by the University of Michigan Committee on Use and Care of Animals. All experiments were conducted using adult rats (300-400 g) maintained on a light: dark cycle of 12: 12 hour (lights on at 6:00 am) and provided with ad libitum food and water.

### Electrode implantation and configuration

Rats were implanted with electrodes for EEG signal recording under surgical anesthesia (1.8% (vol/vol) isoflurane). The EEG signals were recorded through screw electrodes implanted bilaterally on the frontal (anteroposterior (AP): + 3.0 mm; mediolateral (ML): ± 2.5 mm, bregma), parietal (AP: −3.0 mm; ML ± 2.5 mm, bregma), and occipital (AP: −8.0 mm; ML: ± 2.5 mm, bregma) cortices. The ECG signals were recorded through flexible and insulated (except at the tip) multi-stranded wires (Cooner Wires, Chatsworth, CA) inserted into subcutaneous muscles flanking the heart. The EEG and ECG electrodes were interfaced with two six-pin pedestals (Plastics One, Roanoke, VA), and the entire assembly was secured on the skull using dental acrylic.

### Signal acquisition and stroke surgeries

Prior to data collection, rats were acclimatized in the recording chamber. EEG and ECG signals were recorded using Grass Model 15LT physiodata amplifier (15A54 Quad amplifiers) system (Astro-Med, Inc., Quincy, MA) interfaced with BIOPAC MP-150 data acquisition unit and AcqKnowledge software (version 4.1.1, BIOPAC systems, Inc., Goleta, CA). The signals were filtered between 0.1 and 300 Hz and sampled at 1,000 Hz. EEG and ECG recordings were initiated consistently at 10:00 am to control for circadian factors. Baseline signals were recorded for 1 hour. At the end of this baseline recording, permanent BCCAL was performed for SHRPS and WKY rats under surgical anesthesia (Ogata et al. 1976). After BCCAL, recording was continued until sudden death for SHRSP rats and for at most 24 hours for WKY rats.

### Analysis of RR interval (RRI), cardiac arrhythmias, and heart rate variability (HRV)

To analyze the RRI, baseline drift correction was first implemented using second-order Butterworth high-pass filtering with a cutoff frequency at 1 Hz (butter.m and filtfilt.m in Matlab Signal Processing Toolbox; MathWorks Inc., Natick, MA). R peaks of ECG signals were then detected using variable threshold method (Kew and Jeong 2011). Specifically, an amplitude threshold in each nonoverlapping 1 second epoch was applied to select the candidates for R peaks, which can be verified only if the RRI value exceeds a predefined threshold. In this study, the interval threshold was selected as half of the median value of the RRI values in the last 1 second epoch. The automatically detected R peaks were manually validated through a custom user interface developed in Matlab (MathWorks Inc., Natick, MA). As the next step, outliers were removed from the validated RRI using threshold and sliding window averaged filter. RRI was then interpolated using linear interpolation for every 1 minute and plotted for SHRSP and WKY rats (Figure 1B). The mean and standard deviation of RRI during baseline (1 hour before BCCAL), first hour (1st hour after BCCAL), and last hour (1.5-0.5 hour before death) were calculated for all SHRPS (n=9) and WKY (n=8) rats (Figure 1C). To analyze the number and types of cardiac arrhythmias, ECG signals were examined and cardiac arrhythmias were manually labeled and counted using a custom user interface developed in Matlab (MathWorks Inc., Natick, MA).

**Figure 1.**
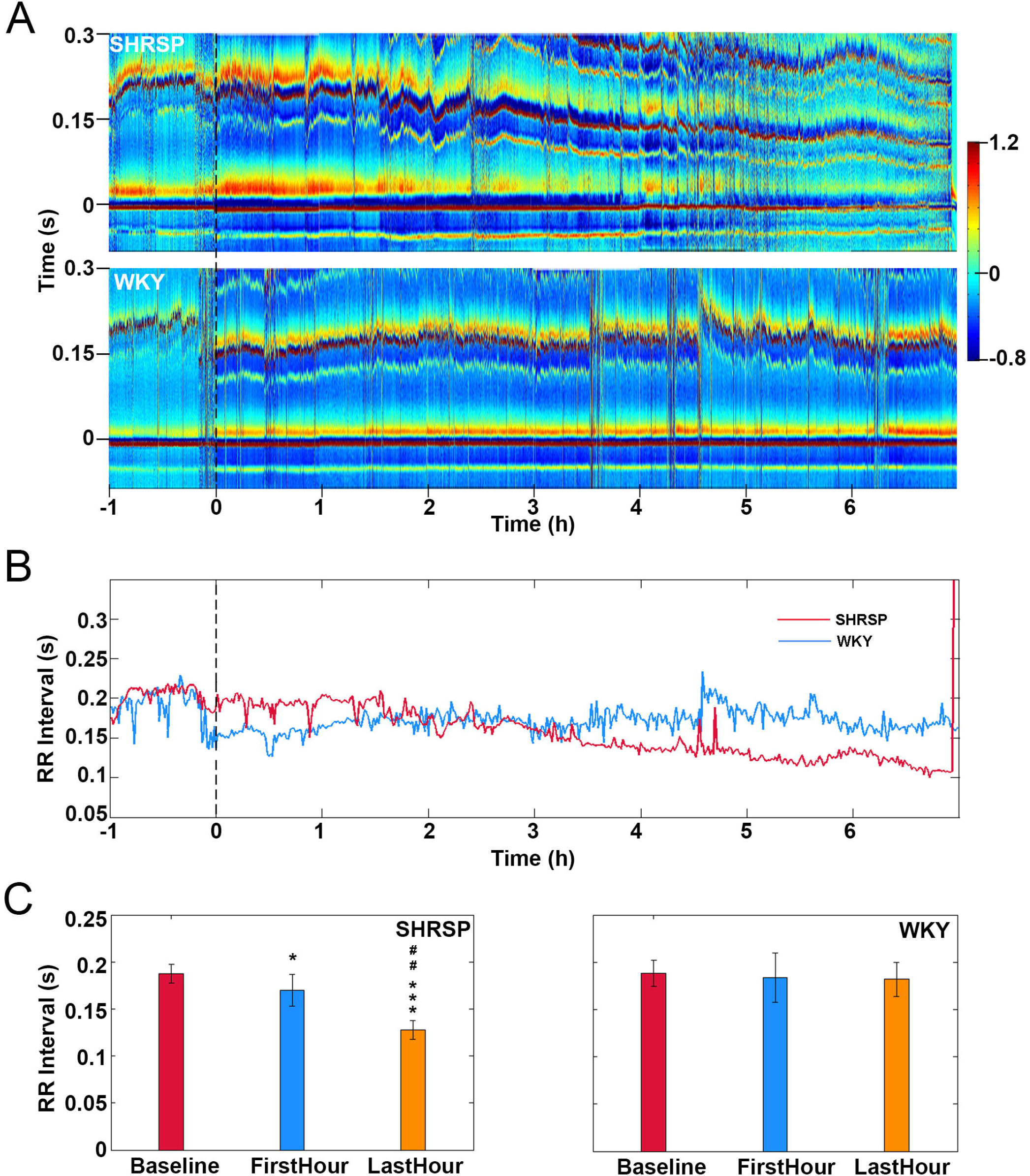
BCCAL results in marked reduction of RRI in SHRSP rats. **(A)** Electrocardiomatrix (ECM) display of ECG signals before and after BCCAL (time 0 sec) for representative rats. **(B)** RRI before and after BCCAL for representative rats. **(C)** RRI during baseline (−1-0 hour), first hour (0-1 hour), and last hour (1.5-0.5 hour before death) for all SHRSP (n=9) and WKY (n=8) rats. Data expressed as mean±SD. *Significant differences over baseline, ##significant differences between first and last hour (*/#p < 0.05, **/##p < 0.01, ***/###p < 0.001).

HRV was analyzed to study the interactions between the sympathetic and parasympathetic nervous system. While high frequency (HF) is considered to reflect parasympathetic activity, low frequency (LF) is thought to be affected by both the sympathetic and parasympathetic activity (Kuwahara et al. 1994). The ratio between LF and HF (LF/HF) reflects the relative balance of sympatho-vagal influences on the heart. Frequency domain analysis of HRV was performed using Fast Fourier transform for LF (0.25-0.8 Hz) and HF (0.8-3 Hz). Frequency selection was based on previous works in rats (Fauchier et al. 2006; Barbosa et al. 2013). The power spectrum of HRV was calculated and expressed in log scale for SHRSP and WKY rats (Figure 1A). The mean and standard deviation of power spectral density for LF, HF, and LF/HF during baseline (1 hour before BCCAL), first hour (1st hour after BCCAL), and last hour (1.5-0.5 hour before death) were calculated for all SHRPS (n=9) and WKY rats (n=8) (Figure 1B).

### Construction of electrocardiomatrix (ECM)

The ECM is designed to facilitate the visualization of RRI, the amplitude, and the morphology of ECG signals. For construction of ECM (Li et al. 2015b), a window centered on the detected ECG R peaks (for example, from −0.1 second to 0.3 second, with 0 corresponding to the time of R-peak) was extracted from the ECG signal after baseline drift correction. All ECG windows were sorted according to the order of R-peak time and then plotted as parallel colored lines to form a colored rectangular image. The intensity of ECG signal was denoted on z-axis, with warmer color indicates positive peaks with higher voltage, while cooler color indicates negative peaks with lower voltage. The color scheme could be adjusted according to the need.

### Analysis of EEG power

The original sampling frequency of 1,000 Hz was first down-sampled to 500 Hz to reduce computing time. A notch filter was used to remove the 60 Hz artifact and its possible super-harmonics. Then EEG power was analyzed using short time Fourier transform based on discrete Fourier transform with 2-second segment size and 1 second overlapping for each frequency bin (0.5-250 Hz with 0.5 Hz bin size; spectrogram.m in Matlab Signal Processing Toolbox, MathWorks Inc., Natick, MA). Each segment was windowed with a Hamming window. The absolute EEG power was expressed on a log scale for SHRSP and WKY rats (Figure 2A). The absolute EEG power on gamma 1 frequency was summed up and plotted for SHRSP and WKY rats (Figure 2B). The mean and standard deviation of absolute EEG power during baseline (1 hour before BCCAL), first hour (1st hour after BCCAL), and last hour (1.5-0.5 hour before death) was calculated for six frequency bands: delta (0.5-5 Hz), theta (5-10 Hz), alpha (10-15 Hz), beta (15-25 Hz), gamma 1 (25-55 Hz), and gamma 2 (65-115 Hz) for all SHRPS (n=9) and WKY (n=8) rats (Figure 2C).

**Figure 2.**
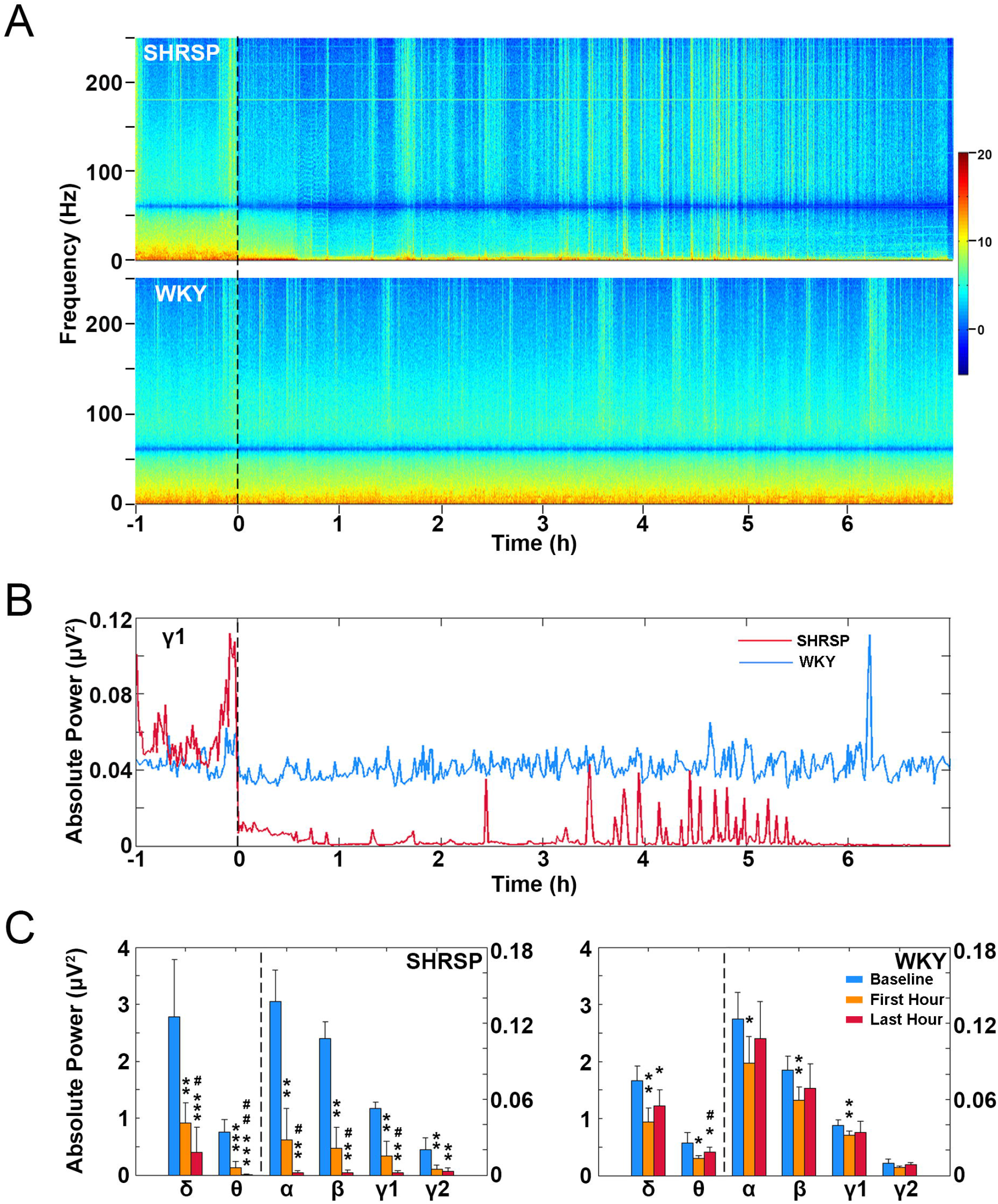
BCCAL results in marked reduction of EEG power in SHRSP rats. **(A)** EEG power spectrum before and after BCCAL (at time 0) for all frequencies in representative rats. **(B)** EEG power before and after BCCAL at gamma 1 frequency in representative rats. **(C)** EEG power during baseline (−1-0 hour), first hour (0-1 hour), and last hour (1.5-0.5 hour before death) for all SHRSP (n=9) and WKY (n=8) rats. Data expressed as mean±SD. *Significant differences over baseline, ##significant differences between first and last hour (*/#*p* < 0.05, **/##*p* < 0.01, ***/###*p* < 0.001).

### Analysis of CCoh and CCCoh

The coherence between 6 EEG channels (CCoh), or between 1 ECG and 6 EEG channels (CCCoh) was measured by amplitude squared coherence (*C_xy_*(*f*)) (mscohere.m in Matlab Signal Processing Toolbox, MathWorks Inc., Natick, MA), which is a coherence estimate of the input signals x and y using Welch’s averaged, modified periodogram method. The magnitude squared coherence estimate (*C_xy_*(*f*) is a function of frequency with values between 0 and 1 that indicates how well signal x corresponds to signal y at each frequency:

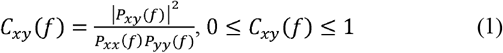

where *P_xx_*(*f*) and *P_yy_*(*f*) are the power spectral density of x and y, and *P_xx_*(*f*) is the cross power spectral density.

In current study, a notch filter was used to remove the 60 Hz artifact and its possible super-harmonics. EEG or ECG signals were then segmented into 2-second epochs with 1-second overlap. The magnitude squared coherence was calculated at each epoch and frequency bin (0.5-250 Hz with 0.5 Hz bin size). The mean coherence between 6 EEG channels (Figure 3A) and among 1 ECG and 6 EEG channels (Figure 4A) were calculated and plotted for frequencies from 0.5 to 250 Hz for SHRSP and WKY rats. The mean CCoh (Figure 3B) and CCCoh (Figure 4B) for gamma 1 frequency band were also plotted for SHRSP and WKY rats. The mean and the standard deviation of coherence among 15 pairs of 6 EEG channels during baseline (1 hour before BCCAL), first hour (1st hour after BCCAL), and last hour (1.5-0.5 hour before death) were calculated for frequencies from delta to gamma 2 for all SHRPS (n=9) and WKY (n=8) rats (Figure 3C). The mean and the standard deviation of percent changes on the amplitude of CCCoh over baseline were calculated for each hour after BCCAL and compared between SHRSP (n=9) and WKY (n=8) rats (Figure 4C, left panel). The duration of epochs with CCCoh 2 times higher than baseline was summed up for each rat and expressed as a percentage of the total duration for that rat. Comparison was then made between SHRSP (n=9) and WKY (n=8) rats for all 6 frequencies (from delta to gamma 2) (Figure 4C, right panel).

**Figure 3.**
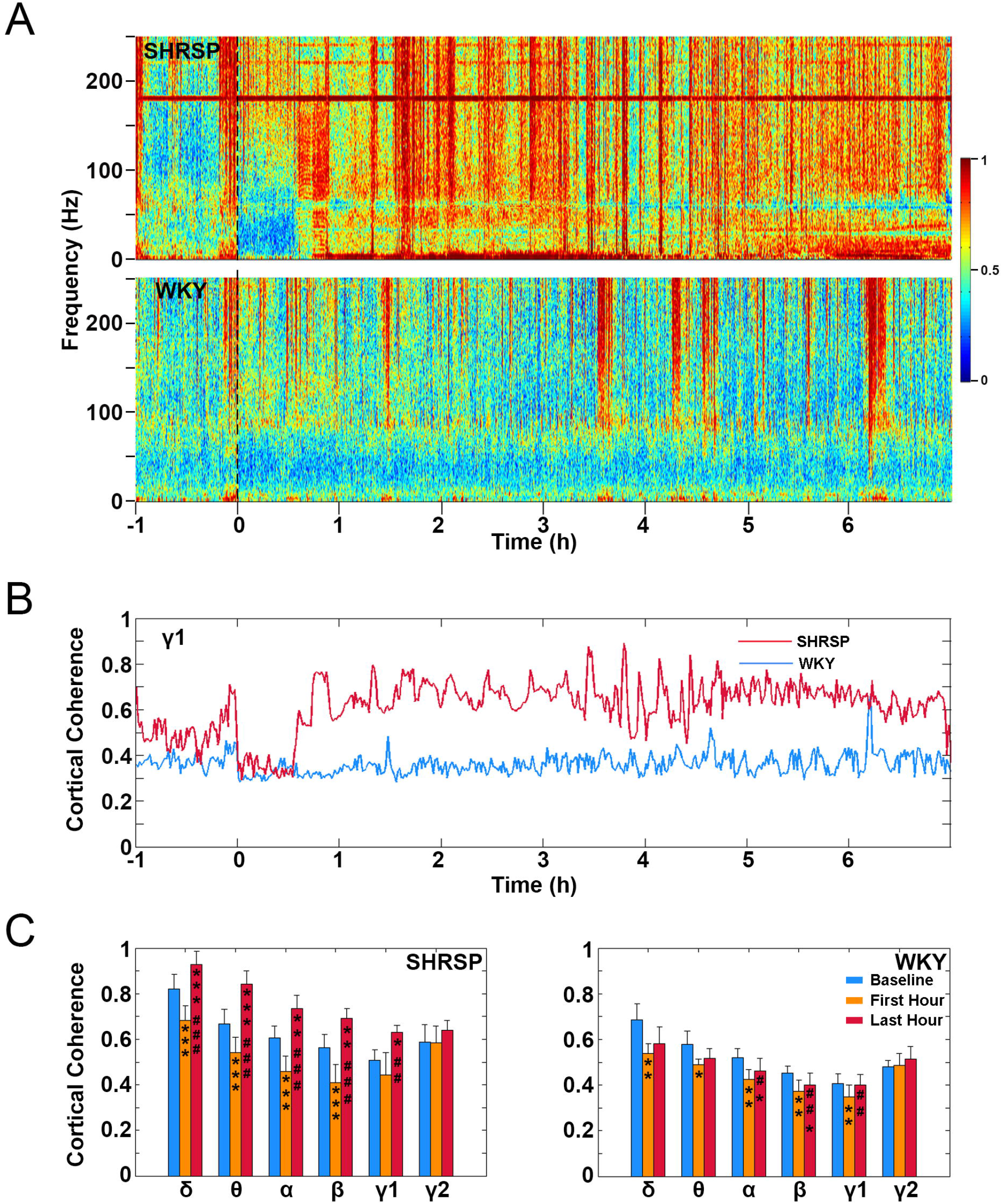
BCCAL results in marked increase of CCoh in SHRSP rats. **(A)** CCoh before and after BCCAL (at time 0) for all frequencies in representative SHRSP and WKY rats. **(B)** CCoh before and after BCCAL for gamma 1 frequency in representative SHRSP and WKY rats. **(C)** CCoh during baseline (−1-0 hour), first hour (0-1 hour), and last hour (0.5-1.5 hour before death) all SHRSP (n=9) and WKY (n=8) rats. Data expressed as mean±SD. *Significant differences over baseline, ##significant differences between first and last hour (*/#*p* < 0.05, **/##*p* < 0.01, ***/###*p* < 0.001).

**Figure 4.**
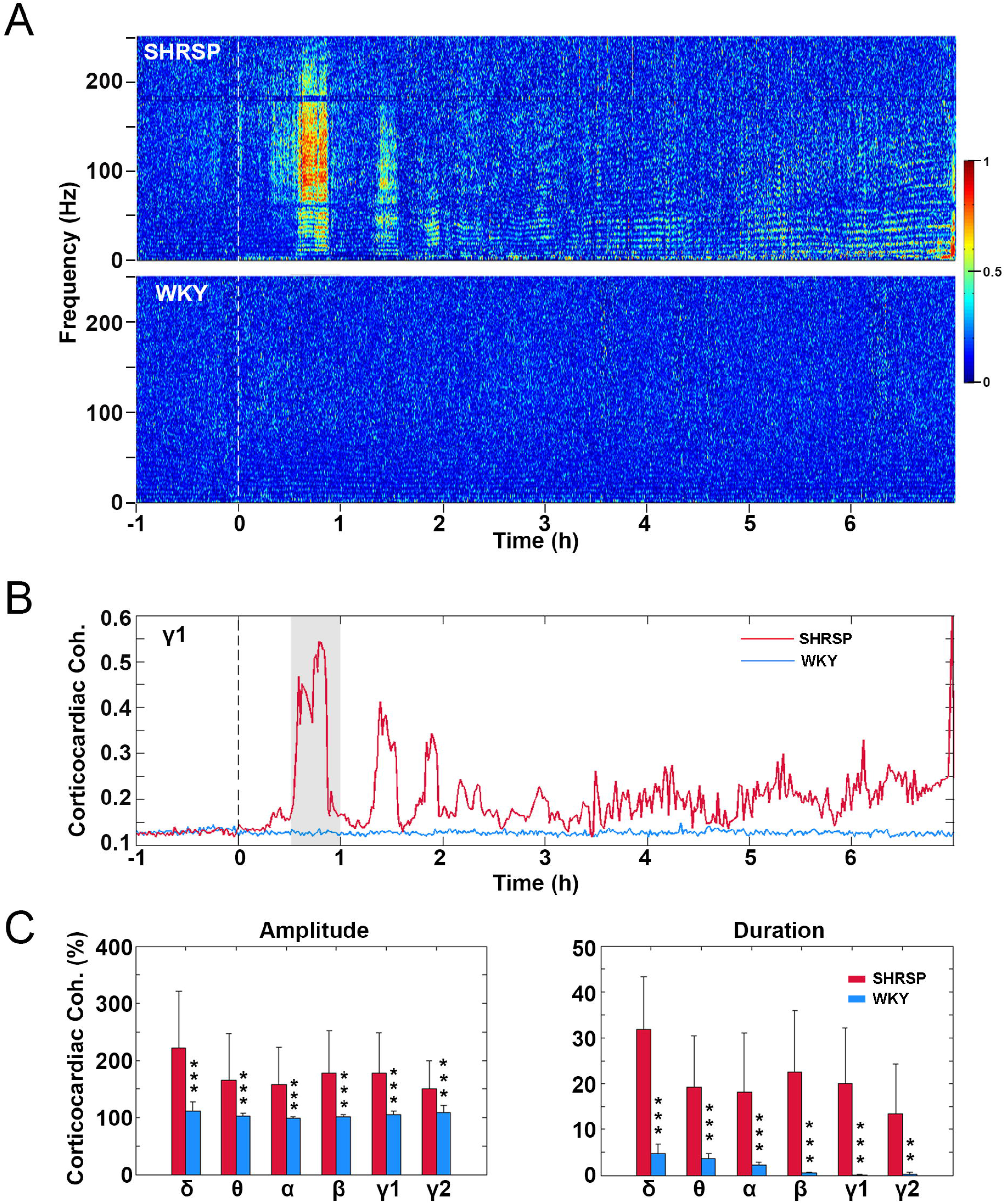
BCCAL results in intermittent surge of CCCoh in SHRSP rats. **(A)** CCCoh before and after BCCAL (at time 0) for all frequencies in representative SHRSP and WKY rats. **(B)** CCCoh before and after BCCAL at gamma 1 frequency in representative rats. **(C)** Percent changes on the amplitude of CCCoh over baseline between all SHRSP (n=9) and WKY (n=8) rats (left panel), percent of signal duration with 2 times higher than baseline CCCoh between all SHRSP (n=9) and WKY (n=8) rats (right panel). Data expressed as mean±SD. *Significant differences between SHRSP and WKY rats (\**p* < 0.05, \*\**p* < 0.01, \*\*\**p* < 0.001).

### Analysis of CCCon

The directional connectivity between EEG and ECG signals was measured by modified (Li et al. 2015a) Normalized Symbolic Transfer Entropy (NSTE) (Lee et al. 2009), which is a nonlinear and model-free estimation of directional functional connection based on information theory. STE measures the amount of information provided by the additional knowledge from the past of the source signal *X*(*X^P^*) in the model describing the information between the past *Y*(*Y^p^*) and the future *Y*(*Y^F^*) of the target signal *Y*, which is defined as following:

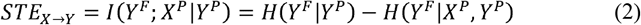

where *H*(*Y^F^|Y^P^*) is the entropy of the process *Y^F^* conditional on its past. Each vector for *Y^F^*, *X^P^* and *Y^p^* is a symbolized vector point. The potential bias of STE was removed with a shuffled data, and the unbiased STE is normalized as follows:

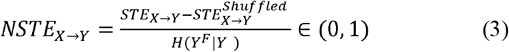

where 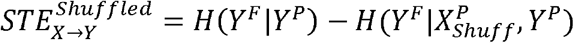. 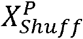is a shuffled data created by dividing the data into sections and rearranging them at random. Therefore, NSTE is normalized STE (dimensionless), in which the bias of STE is subtracted from the original STE and then divided by the entropy within the target signal, *H*(*Y^F^|Y^P^*).

For CCCon, the feedback connectivity 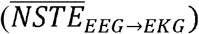 was calculated by averaging NSTE over 6 pairs of EEG channels to ECG channel, which are defined as follows:

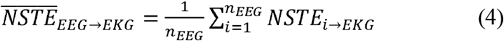

where *n_EEG_* = 6. The feedforward connectivity 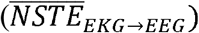 from the ECG to 6 EEG channels is vice versa.

Specifically, we first filtered EEG and ECG signals into 6 frequency bands (from delta to gamma 2), and then segmented the filtered signals into 2-second long epochs with 1-second overlapping. The mean CCCon 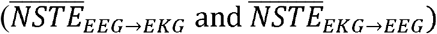 were sequentially calculated for each epoch and each frequency band. Three parameters: embedding dimension (*d_E_*), time delay (τ), and prediction time (*δ*), were required in the calculation. In this study, we selected the parameter setting that could yield maximum *NSTE_X→Y_* by fixing the embedding dimension (*d_E_*) at 3, and optimizing prediction time *δ* (from 1 to 50, corresponding to 1-50 ms with the sampling frequency of 1,000 Hz) and time delay *τ* (1-300 ms). The same procedure was used to calculate *NSTE_Y→X_*, provided that the information between two signals is transferred through different neuronal pathway. The feedback and feedforward CCCon were plotted for theta and gamma 1 frequency for a sample epoch that shows a surge of CCCoh (Figure 5A). The mean and standard deviation of CCCon epochs during baseline (1 hour before BCCAL), first hour (1st hour after BCCAL), and last hour (1.5-0.5 hour before death) was calculated for six frequency bands for all SHRPS (n=9) rats (Figure 5B).

**Figure 5.**
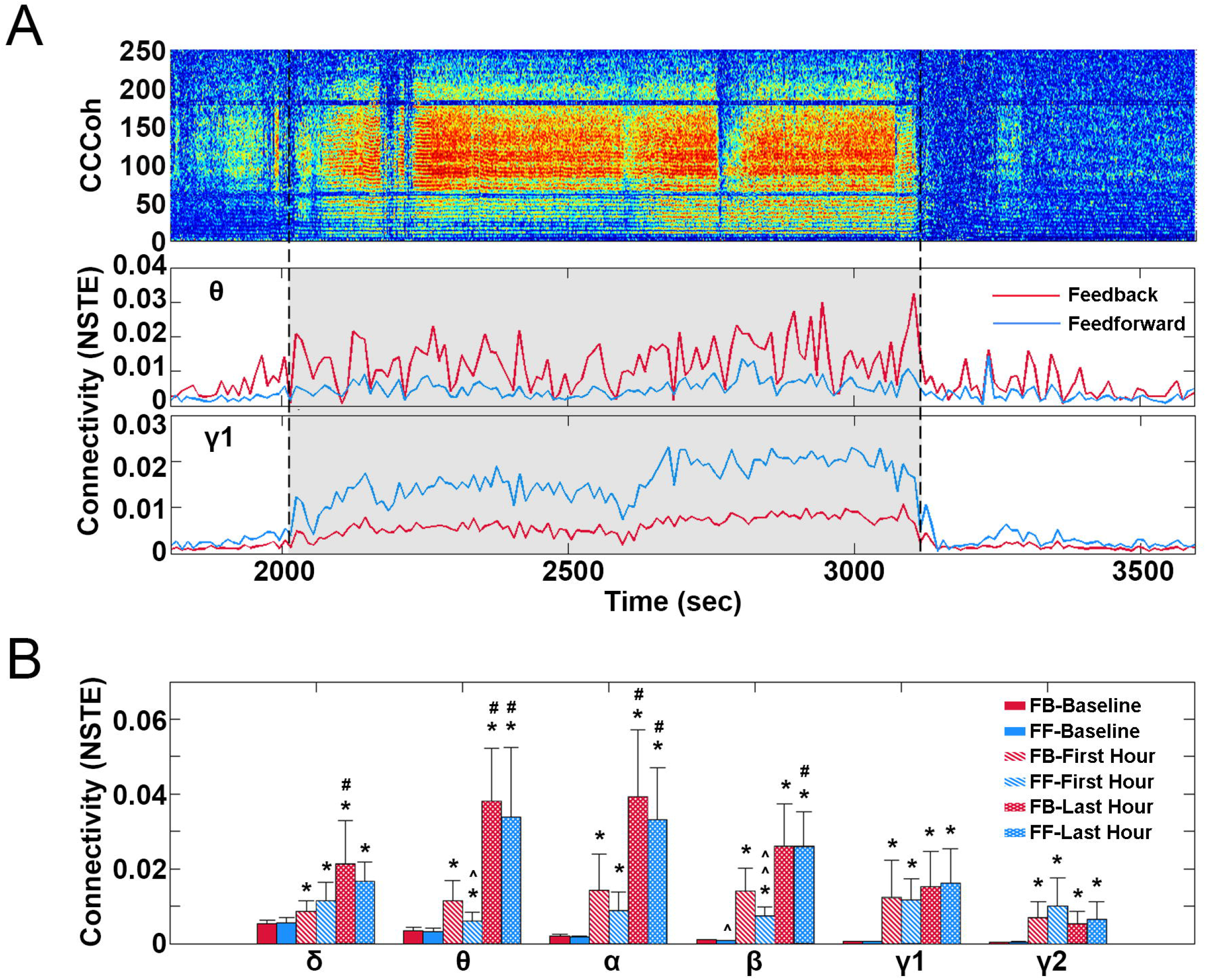
BCCAL results in increase of CCCon in SHRSP rats. **(A)** CCCon at theta and gamma 1 frequencies for one representative epoch that shows high CCCoh. **(B)** CCCon for selected epochs during baseline (−1-0 hour), first hour (0-1 hour or first few hours after BCCAL), and last hour (0-1 hour before sudden death). Data expressed as mean±SD. *Significant differences over baseline, #significant differences between first and last hour, ^significant differences between feedforward and feedback CCCon (*/#/^p < 0.05, **/##/^^p < 0.01, ***/###/^^^p < 0.001).

### Statistical analysis

For all the statistical analyses, Shapiro-Wilk normality test was first implemented to determine if the data was normally distributed. To test the differences of RRI (Figure 1C), EEG power (Figure 2C), CCoh (Figure 3C), CCCon (Figure 5B), and HRV (Figure 6C) among baseline, first hour, and last hour, repeated measures ANOVA with post hoc Pair-Sample T-test (for normally distributed data) or Friedman Test with Wilcoxon post hoc comparisons (for non-normally distributed data) were conducted. To analyze the differences of CCCoh between SHRSP and WKY (Figure 4C), and the differences between feedback and feedforward CCCon (Figure 5B), independent-sample T-test (for normally distributed data) or Mann-Whitney (for non-normally distributed data) test was used. For all the comparisons, *p* < 0.05 was considered as statistically significant. Statistical analyses were performed using the software SPSS (version 19.0; IBM SPSS Statistics).

**Figure 6.**
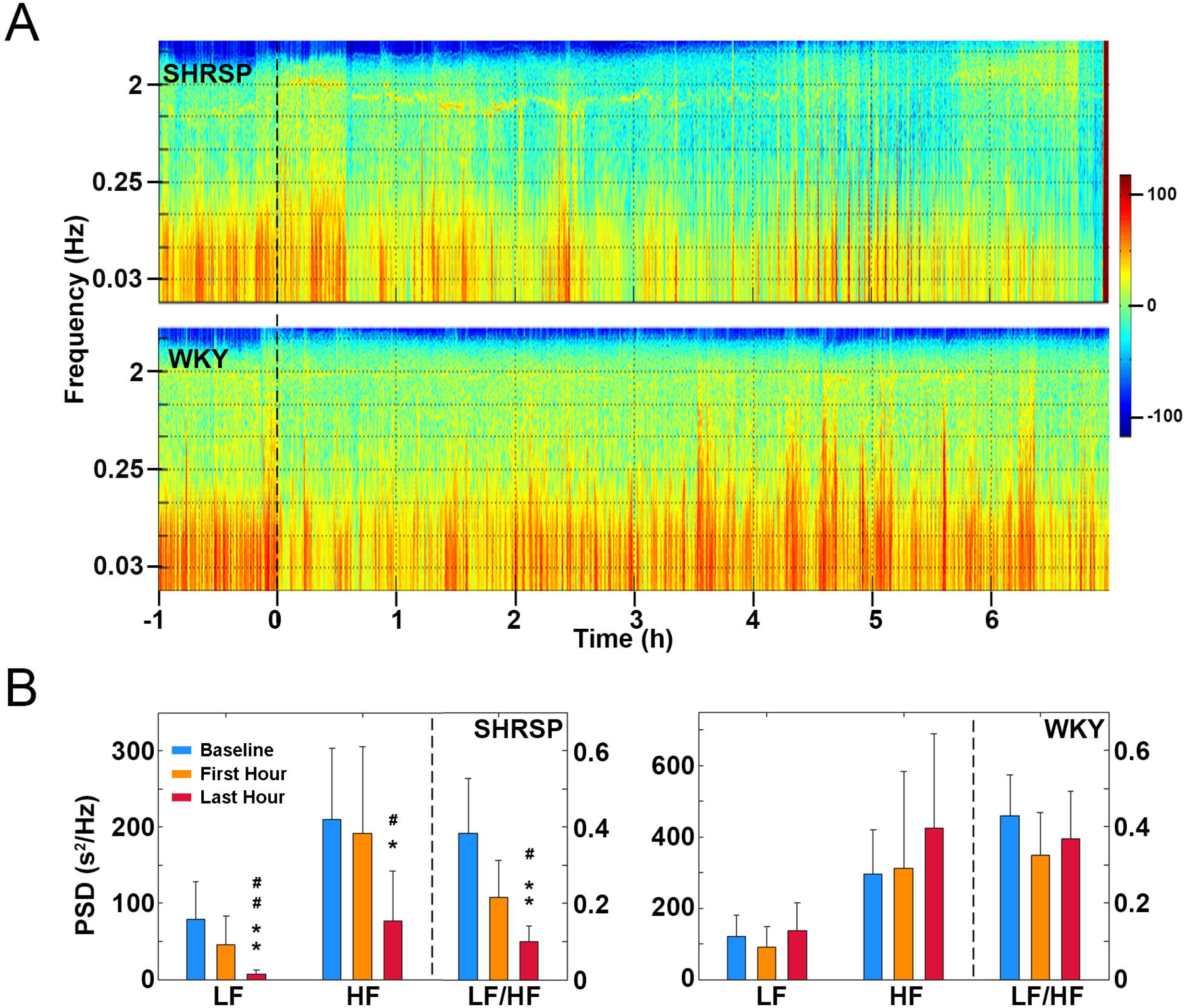
BCCAL results in marked reduction of HRV for SHRSP rats. **(A)** HRV before and after BCCAL (at time 0) for representative SHRSP and WKY rats. z axis represents power spectral density (PSD). Warmer color represents stronger PSD. **(B)** PSD of low frequency (LF), high frequency (HF), and LF/HF during baseline (−1-0 hour), first hour (0-1 hour), and last hour (0.5-1.5 hour before death) for all SHRSP (n=9) and WKY (n=8) rats. Data expressed as mean±SD. *Significant differences over baseline, #significant differences between first and last hour (*p < 0.05, **p < 0.01, ***p < 0.001).

## Results

### Forebrain ischemia claimed 100% mortality in SHRSP rats within 14 hours

A total of 9 SHRSP and 8 WKY rats underwent BCCAL procedure, which caused 100% mortality in SHRSP rats within 14 hours. In contrast, no mortality was observed in WKY control rats. The survival length of SHRSP rats ranged from 2.89 hours to 13.92 hours, with a mean length of 8.13 hours.

### SHRSP rats suffering from forebrain ischemia exhibited a marked decrease in RRI and increase in cardiac arrhythmias

The effects of forebrain ischemia on cardiac function was investigated. ECM was first generated to display dynamic changes of RRI and cardiac arrhythmias before and after BCCAL procedure in representative SHRSP and WKY rats (Figure 1A). The temporal reduction of RRI (i.e., increase of heart rate) is clearly seen in the SHRSP rat (Figure 1B). During baseline (−1 to 0 hour) condition, SHRSP and WKY rats displayed similar RRI of about 0.2 seconds [heart rate of 300 beats/minute (bpm)). After BCCAL, RRI for SHRSP rat exhibited gradual and persistent decline until the sudden collapse of cardiac function at 7^th^ hour after ischemic stroke. In sharp contrast, RRI for the WKY rat showed visible reduction from 0.2 s to 0.15 s, 10 min prior to the initiation of the ischemic procedure, which persisted during the entire process (Figure 1B). These features are conserved for all SHRSP and WKY rats (Figure 1C). As shown in the left panel of Figure 1C, RRI for SHRSP rats was significantly lower in first hour after ischemia (0.17±0.02 second (353 bpm)) than the baseline (0.19±0.01 second (316 bpm)). Moreover, SHRSP rats showed a further significant reduction of RRI in the last hour (1.5-0.5 hour before sudden death) (0.13±0.01 second (462 bpm)) compared to baseline and first hour after BCCAL. However, no significant differences on RRI were found for WKY rats among baseline, first hour after BCCAL, and last hour before sudden death (right panel in Figure 1C).

Cardiac arrhythmias before and after forebrain ischemia are summarized in Table 1. A total of 9 common arrhythmias were identified. Among them, the average hourly occurrence of premature ventricular contraction (PVC), sinus pause (SP), junctional rhythm (JR), and second-degree heart block (2HB) significantly increased after ischemic stroke for both SHRSP and WKY rats. However, significant increase in average hourly occurrence for premature atria contraction (PAC), third-degree heart block (3HB), junctional escape beat (JEB), ventricular escape beat (VEB), and marked sinus bradycardia (MSB) over baseline was only observed in SHRSP rats. In addition, during the last 30 minutes (ending phase) before death, agonal signals including second-degree heart block, third-degree heart block, junctional escape beat, ventricular escape beat, and marked sinus bradycardia remarkably increased for SHRSP rats, but were not identified in any of the WKY rats at comparable times after forebrain ischemia.

**Table 1.**
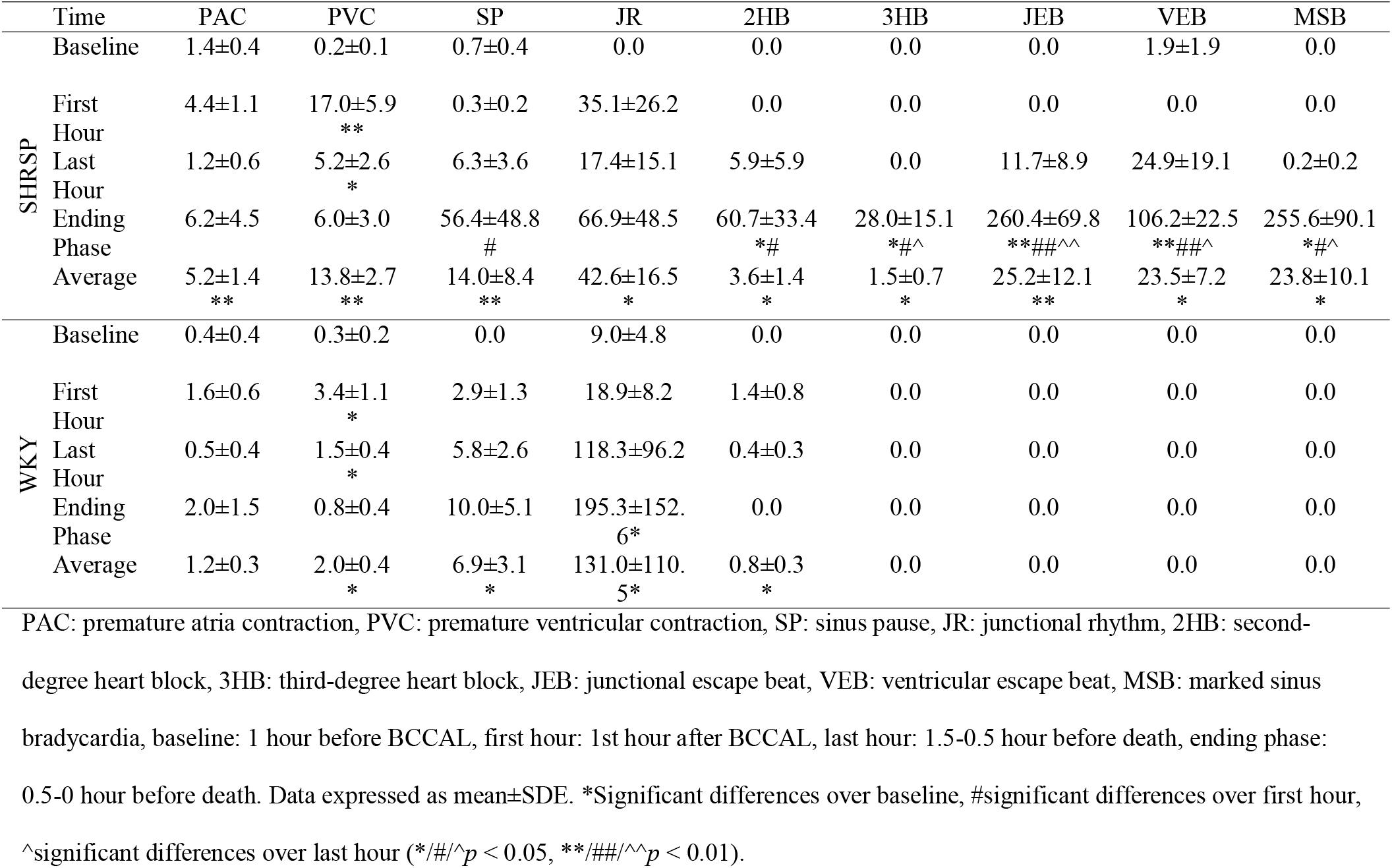
Hourly occurrence of cardiac arrhythmias per rat.

### SHRSP rats exhibited a marked reduction of EEG power following forebrain ischemia

To explore the influence of forebrain ischemia on the electrical activity of the brain, average EEG power across all 6 cortical channels before and after BCCAL was analyzed. Figure 2A shows the EEG power spectrum for all frequencies (0-250 Hz) in representative SHRSP (upper panel) and WKY (lower panel) rats. As shown in the figure, EEG power was dramatically reduced for SHRSP rat immediately after BCCAL and maintained for all frequency bands at the low levels until sudden death (upper panel in Figure 2A). However, EEG power for WKY rat did not exhibit obvious changes over the entire course (lower panel in Figure 2A). The temporal changes of EEG power at gamma 1 frequency (Figure 2B) illustrates how EEG power diminishes after BCCAL specifically for SHRSP, but not WKY, rats. In contrast to the nearly stable EEG power in control WKY rat, EEG power for SHRSP rat demonstrated a dramatic reduction immediately following forebrain ischemia, which then continued to decline until sudden death. Statistical analyses were conducted to compare the changes of EEG power among baseline, first hour after BCCAL, and last hour (0.5-1.5 hour before sudden death) in all SHRSP and WKY rats. As shown in Figure 2C, the significant reduction of EEG power in both first hour after BCCAL and last hour before sudden death compared to baseline was found consistently for all SHRSP rats across all 6 frequencies, which include delta, theta, alpha, beta, gamma 1, and gamma 2 (left panel in Figure 2C). Additionally, the EEG power for last hour was also significantly lower than first hour for delta, theta, alpha, beta, and gamma 1 frequencies. In contrast, while showing significant reduction of EEG power in the first hour after forebrain ischemic stroke in most frequencies (delta, theta, alpha, beta, and gamma 1), WKY rats demonstrated a rebound increase of EEG power in the last hour for all frequencies (right panel in Figure 2C).

### SHRSP rats exhibited a significant increase of CCoh following forebrain ischemia

To explore the influence of forebrain ischemia on functional connectivity (coherence) of the brain, CCoh before and after BCCAL was analyzed. Figure 3A shows the CCoh before and after BCCAL for all frequencies (up to 250 Hz) in representative SHRSP and WKY rats. As shown in the figure, there is a marked decrease of CCoh at low frequencies (0-55 Hz) and increase of CCoh for high frequencies (65-250 Hz) within 1 hour after BCCAL in SHRSP rat (upper panel in Figure 3A). After this early phase reduction of low frequency CCoh, CCoh for all frequencies demonstrated a dramatic and sustained increase until sudden death. However, for WKY rats, while the CCoh at high frequency (65-250 Hz) showed sporadic increases after BCCAL, the CCoh for low frequency bands (0-55 Hz) decreased after ischemic stroke (lower panel in Figure 3A). Figure 3B displays the CCoh at gamma 1 frequency for the same pair of SHRSP and WKY rats. In SHRSP rat, the CCoh for gamma 1 frequency showed a marked reduction within the first hour of ischemia, which immediately increased to values that are greater than the baseline and persisted for the rest of the recording. In contrast, WKY rat showed a small reduction of CCoh after BCCAL and remained at low values for the remaining of the ischemic period. Statistical analyses were conducted to compare the CCoh changes among baseline, first hour after BCCAL, and last hour (0.5-1.5 hour before sudden death) for both SHRSP and WKY rats. As shown in Figure 3C, in both SHRSP (left panel) and WKY (right panel) rats, there was a dramatic reduction of CCoh in first hour after forebrain ischemic stroke than baseline for 5 frequencies (delta to gamma 1). In last hour, however, CCoh increased to a level that was significantly greater than baseline for SHRSP rats (delta to gamma 1), whereas in WKY rats, CCoh in last hour was recovered to a level that is higher than first hour but lower than baseline level.

### Intermittent surge of functional connectivity between the heart and the brain in SHRSP rats following forebrain ischemia

To investigate the impact of forebrain ischemia on the electrical signal synchronization between the brain and the heart, CCCoh before and after BCCAL was calculated. Figure 4A shows the CCCoh for all frequencies in representative SHRSP and WKY rats. Intermittent surge of CCCoh was observed for SHRSP rats after BCCAL (upper panel in Figure 4A). However, the CCCoh for WKY rats was nearly undetectable in both baseline and after BCCAL procedure (lower panel in Figure 4A). Figure 4B displays the CCCoh at gamma 1 frequency for the same pair of SHRSP and WKY rats. In contrast to WKY rat, which demonstrated no obvious changes in CCCoh before and after BCCAL, SHRSP rat showed a dramatic surge of CCCoh at gamma 1 frequency within the first hour of ischemic injury, and a continued and intermittent surge of CCCoh throughout the remainder of the recorded period. Statistical analyses were conducted to compare the changes on the amplitude of CCCoh and the duration of high CCCoh epochs after BCCAL between SHRPS and WKY rats (Figure 4C). As shown in figure, a large increase in the amplitude of CCCoh after BCCAL over baseline (0.50-1.21-fold increase) was identified for SHRSP rats for all frequencies, but was not observed for WKY rats, as indicated by the 0.01-0.11-fold change of CCCoh amplitude before and after BCCAL among all frequencies (left panel in Figure 4C). Statistical analysis suggests that the increase of CCCoh amplitude after BCCAL was significantly higher in SHRSP than WKY rats. Right panel shows the percentage of signal duration with CCCoh amplitude 2 times higher than baseline level over the total duration of the signal in all SHRSP and WKY rats. While 13-32% of signals after BCCAL had CCCoh that is 2 times higher than baseline in SHRSP rats, the CCCoh for WKY maintain at low level after BCCAL, with less than 5% of the signals that has CCCoh that is 2 times higher than baseline. Statistical analysis suggests that the percentage of CCCoh epochs that are 2 times larger than baseline after BCCAL was significantly larger in SHRSP than WKY rats (right panel in Figure 4C).

### Intermittent surge of directional connectivity between the heart and the brain in SHRSP rats following forebrain ischemia

To examine the impact of forebrain ischemia on the directional communication between the brain and the heart, CCCon before and after forebrain ischemia was analyzed. Figure 5A shows the feedback (from the brain to the heart or efferent) and feedforward (from the heart to brain or afferent) CCCon in a sample epoch that has high CCCoh (corresponding to the shaded region in Figure 4A) within the first hour after BCCAL in representative SHRSP rat. As shown in the figure, a marked surge of connectivity in both theta and gamma 1 frequencies was detected in feedforward as well as feedback directions when CCCoh was elevated. However, when the CCCoh was low, the heart-brain connectivity was low in both directions for both theta and gamma 1 frequencies. Statistical analyses were conducted to compare the changes of feedforward and feedback CCCoh for all frequencies among epochs selected from baseline, first hour (0-1 hour or first few hours after BCCAL), and last hour (0-0.5 hour before sudden death) (Figure 5B). As shown in the figure, the significant increase of CCCon after forebrain ischemia over baseline values was found in all SHRSP rats, both immediately following ischemia (first hour) or during near-death stage (last hour), for all tested frequencies. Additional increase at the late stage of ischemia (last hour) over the early stage of ischemia (first hour) was observed for delta (feedback), theta (both feedback and feedforward), alpha (both feedback and feedforward), and beta (feedforward) frequencies. Directional asymmetry was also detected for theta and beta frequencies at the early stage of ischemia, with feedback connectivity dominated over feedforward connectivity.

### SHRSP rats displayed a marked reduction of HRV following forebrain ischemia

To understand the impact of forebrain ischemia on autonomic regulation of cardiac function during ischemic stroke-induced sudden cardiac arrest, HRV analysis was conducted for all the rats. The HRV before and after BCCAL in representative SHRSP and WKY rats was shown in Figure 6A. In SHRSP rat, HRV exhibited reduction at both frequency domain and time domain, showing precipitous decline especially at its last hour (upper panel in Figure 6A). In contrast, WKY rat showed an early reduction of HRV in first and second hour following BCCAL procedure, which was then recovered to normal levels in the following hours (lower panel in Figure 6A). These changes of HRV are conserved in all SHRSP and WKY rats. As shown in Figure 6B, in SHRSP rats, both low frequency (LF, 0.25-0.8 Hz) and high frequency (HF, 0.8-3 Hz) components as well as the ratio of low and high frequency (LF/HF) displayed a reduction in first hour after BCCAL and last hour (0.5-1.5 hour before sudden death) than baseline condition (left panel in Figure 6B). Remarkably, the LF, HF, and LF/HF at last hour are significant lower compared to both baseline and first hour. In contrast, WKY rats showed an initial decline of LF and LF/HF in first hour after BCCAL, which was followed by a recovery of LF, HF, and LF/HF in last hour before sudden death. Different from SHRSP rats, these changes of HRV in WKY rats are not significant (right panel in Figure 6B).

## Discussion

This is the first study to examine the dynamic changes of the brain, the heart, and the functional interactions between the brain and the heart in forebrain ischemic stroke-induced sudden cardiac arrest rat models. Our data demonstrated that there is a dramatic decline of cardiac functionality (decrease of RRI and increase of cardiac arrhythmias), disruption of the autonomic nervous system (decrease of HRV), and reduction of cortical electrical activity (EEG power) during forebrain ischemic cardiac arrest. Importantly, we identified a global increase of functional synchronization within the brain (CCoh), as well as a marked and intermittent surge of brain-heart coupling, indicated by high levels of CCCoh and CCCon, from the onset of ischemia until death in rats that suffered from sudden cardiac arrest. These results suggest that abnormal brain-heart connection may be the mechanism for forebrain ischemic stroke-induced sudden cardiac arrest and the surge of CCCoh may be used as a biomarker to predict the risk of sudden death.

### Animal model for sudden cardiac arrest

Unlike our previous studies (Borjigin et al. 2013; Li et al. 2015a; Tian et al. 2018) where rats experienced global hypoxia (asphyxia) that affects the brain and the heart simultaneously, in the current model, we investigated brain-heart connection in sudden cardiac arrest induced by forebrain ischemic stroke. Comparatively speaking, forebrain ischemia model is clinically more impactful than asphyxia model because it tests the direct influence of brain ischemia on the function of the brain and the heart. In asphyxia models, the cortex, brainstem, spinal cord, and the heart are globally affected by the experimental insult, making it difficult to test the role of the brain during the dying process. BCCAL in SHRSP rats is a well-established model for forebrain ischemia (Katayama et al. 1984; Lobanava et al. 2008). One previous study showed that occlusion of the bilateral carotid artery caused the death of SHRSP rats within 6 hours, whereas the control WKY rats died within 8 hours after the occlusion (Kakihana and Nagaoka 1983). In our experiment, BCCAL induced 100% mortality for SHRSP rats within 14 hours. However, no mortality was observed in WKY rats. The high mortality in previous study compared to our study may due to the genetic variations for outbred animals, as well as the high salt Japanese diet (4% sodium chloride in Japanese diet vs. 0.4% sodium chloride in standard chow), which accelerates the development of hypertension in SHRSP rats (Matsuo and Nagaoka 1981; Stier et al. 1988; Schmidlin et al. 2005). Nevertheless, large similarities in mitochondria abnormalities and cerebral blood vessel deficits during the development of stroke have been identified between SHRSP rats and humans, making BCCAL in SHRSP rats one of the more relevant model for studying sudden cardiac arrest induced by forebrain cerebral ischemia (Lobanava et al. 2008).

### Deterioration of cardiac function and autonomic nervous system functionality during forebrain ischemic stroke-induced sudden cardiac arrest

Cardiac arrhythmias and ECG abnormalities are commonly observed after acute cerebrovascular events, such as ischemic stroke, intracerebral hemorrhage, and subarachnoid hemorrhage (Goldstein 1979; Daniele et al. 2002). In stroke patients, cardiac arrhythmias, especially the malignant ventricular arrhythmias triggered by the impairment of the central autonomic nervous system structures and catecholamine storm, are highly prevalent (Myers et al. 1982; Mäkikallio et al. 2004). It is also known that ventricular tachycardia (VT), ventricular fibrillation (VF), and asystole are often the fatal cardiac arrhythmias occurring right before sudden death (Bayes de Luna et al. 1989). However, none of previous studies has characterized the number and types of cardiac arrhythmias leading to cardiac arrest in forebrain ischemic stroke rat model, except for one earlier study which reported the disturbance of cardiac rhythm and increase in the number of PVC in SHRSP rats after BCCAL (Kakihana et al. 1983). In our study, we showed that common arrhythmias including PVC, PAC, sinus pause, junctional rhythm and second-degree heart block significantly increased after the surgery in both SHRSP and WKY rats, although on average more arrhythmias were found in SHRSP rats than WKY rats. The occurrences of those arrhythmias are sporadic and irregular in both strains. Importantly, there is a sudden increase of cardiac arrhythmias in last 30 minutes before death only in SHRSP rats. This data suggests that the deterioration of cardiac function is very acute, which further proves the validity of current model for investigating the mechanism of sudden cardiac death. Interestingly, different from asphyxic cardiac arrest model, in which third-degree heart block, junctional escape beat, ventricular escape beat-, ventricular tachycardia, and ventricular fibrillation are the last cardiac arrhythmias before sudden death (Tian et al. 2018), in forebrain ischemic cardiac arrest model, ventricular tachycardia and ventricular fibrillation are not identified. Instead a significant increase of marked sinus bradycardia was observed in SHRPS rats right before sudden death.

HRV has been widely used for studying the autonomous nervous system control of cardiovascular function (Task force et al. 1996). Reduced HRV has been associated with the risk of myocardial infarction (Lombardi et al. 1987; Buchanan et al. 1993), congestive heart failure (Saul et al. 1988), and sudden cardiac death (Politano et al. 2008; Wu et al. 2014). Parallel with previous findings, our study demonstrated a significant reduction of LF, HF, and the ratio of LF/HF in SHRSP rats that suffered from sudden cardiac death. Importantly, the control WKY rats that received the same BCCLA procedure did not display significant changes of HRV during the entire process. These results indicate that forebrain ischemia leads to reduced level of activity for both the sympathetic and parasympathetic nervous system. The reduction of the autonomic nervous system functionality may be the cause for the dramatic increase of marked sinus bradycardia during the final phase of forebrain ischemic leading to death. We reported previously that the sympathetic nervous system is over-activated during asphyxic cardiac arrest, since the blockade of sympathetic nervous system via either spinal cord transection or adrenergic blockers prolongs the electrical activities of both the brain and the heart (Li et al. 2015a; Tian et al. 2018). The cardiac arrhythmias observed in the ending phase in asphyxic rats are therefore VT and VF. The differences of cardiac arrhythmias and autonomic nervous system functionality between global ischemia/hypoxia and focal ischemia indicate that cardiac response to global ischemia (asphyxia) and forebrain ischemia are due to different (autonomic) mechanisms.

### Decrease of cortical power and increase of cortical coherence during forebrain ischemic stroke-induced sudden cardiac arrest

Consistent with previous studies in mice (Weitzel et al. 2016) and rats (Borjigin et al. 2013) that suffered from cardiac arrest induced by injection of potassium chloride, severe and sustained reduction in EEG power was identified in SHRSP rats after forebrain ischemic stroke. As expected, an initial decline and a subsequent recovery of EEG power was found in WKY rats that survived (Figure 2). A previous study in spontaneously hypertensive rats exposed to forebrain ischemia demonstrated that 20-minute BCCAL plus hypotension produces dramatic increase in delta power and decrease in theta, beta, and alpha activities (Mariucci et al. 2003), comparable to what we found (Figure 2). In addition, similar to what we found for SHRSP and WKY rats, EEG activity also recovered to normal values more quickly in WKY rats than in SHR rats, in which alpha and beta power did not recover even at 6 days of reperfusion (Mariucci et al. 2003). The irreversible influence of forebrain ischemic stroke on brain electrical activity in SHRSP and SHR rats, but not WKY rats, may result from the abnormal cerebral vascular structure, hypertension, and altered hemodynamics of spontaneous hypertensive rats, all of which could render SHRSP and SHR rats more susceptible to ischemic insult than normotensive rats (Ogata et al. 1976; Fujishima et al. 1980; Kakihana et al. 1983; Coyle 1986; Brint et al. 1988; Duverger and MacKenzie 1988; Lobanova et al. 2008).

Immediately after BCCAL, the functional synchronization (CCoh) between different cortical regions significantly decreased in all rats in multiple frequencies (Figure 3C). This result was consistent with findings from previous studies in Wistar rats exposed to BCCAL and in patients with acute thalamic ischemic stroke, both of which showed a decrease of cortical coherence or functional connectivity within the brain after ischemia (Kozhechkin et al. 2009; Liu et al. 2016). Interestingly, CCoh in SHRSP rats exhibited a dramatic surge (hyperconnectivity) within one hour of cardiac arrest, indicating that the brain is internally super-activated at near-death. The neurophysiological mechanisms underlying the marked surge of the functional coupling within the brain after ischemia is still unknown. Nevertheless, the large increase of brain functional synchronization after forebrain ischemic stroke until sudden death is consistent with our earlier findings in both potassium chloride injection- and asphyxia-induced cardiac arrest models, in which a dramatic increase of both functional and effective connectivity was identified within the brain before sudden death (Borjigin et al. 2013; Li et al. 2015a). The identification of dramatic surge of CCoh exclusively in rats that suffered from focal ischemic stroke provides further experimental support for our central hypothesis that the brain plays an active role in the dying process.

### Increase of brain-heart connection during forebrain ischemic stroke-induced sudden cardiac arrest

Despite growing evidence that abnormal interaction between the brain and the heart is the major cause of sudden death, the dynamical changes of brain-heart connection have not been characterized in any neurogenic sudden death models due to the lack of effective methods (Samuels 2007; Dorrance and Fink 2015; Gonzales-Portillo et al. 2016). In previous study, we developed novel biomarkers, CCCoh and CCCon, to investigate the synchronization and bidirectional signal communication between the brain and the heart in asphyxic cardiac arrest models (Li et al. 2015a; Tian et al., 2018). Consistent with the earlier study (Li et al. 2015a), a surge of coherence (CCCoh) was identified in SHRSP rats that ultimately died from forebrain ischemia (Figure 4). In addition, strong bi-directional information transfer between the brain and the heart (CCCon) was also found in SHRSP rats whenever there was increased brain-heart coupling (Figure 5). The strong functional and directional corticocardiac connectivity in dying rats observed in the SHRSP rats demonstrate similar mechanisms underlying the sudden death in both models (global asphyxia and focal cerebral ischemia). There are, however, two differences of CCCoh and CCCon between asphyxia- and forebrain ischemic stroke-induced sudden cardiac arrest models: 1) high level of CCCoh and CCCon was detected for all frequencies (0-250 Hz) in forebrain ischemic cardiac arrest model, whereas in asphyxic cardiac arrest model (Li et al., 2015a), CCCoh was clustered at low frequency ranges (0-55 Hz); 2) The CCCoh in SHRSP rats after forebrain ischemic stroke is intermittently distributed, however, in asphyxic cardiac arrest model, CCCoh displays homogeneous and continuous pattern. Additional studies are clearly warranted to investigate the mechanisms of stroke-induced sudden death.

## Conclusion

This study demonstrated that forebrain ischemic stroke stimulates elevated brain-heart electrical signal coupling and communication that are highly associated with the increase of cardiac arrhythmias, disruption of the autonomic nervous system, and the risk of sudden death. It further suggests that the surge of corticocardiac coupling may be used as a potential biomarker to predict sudden death. This study may improve mechanistic understanding of how the ischemic brain interact with the heart leading to sudden cardiac arrest.

## Acknowledgements

We thank Drs. Richard Keep, Daniel Beard, Anuska Andjelkovic-Zochowska, and UnCheol Lee for their helpful discussions and comments, and the Department of Molecular and Integrative Physiology for support.

